# The brown alga *Saccharina japonica* features distinct vanadium-dependent bromoperoxidases and iodoperoxidases

**DOI:** 10.1101/243410

**Authors:** Shan Chi, Tao Liu, Hongxin Yin, Xin Xu, Weiming Zhu, Yi Wang, Cong Wang, Hui Lv

## Abstract

Marine algae have an extraordinary ability to absorb halogens which provide algae with an inorganic antioxidant impacting atmospheric chemistry. Although brown algal Laminariales species are the most efficient iodine accumulators among all living systems, and *Saccharina japonica* is the primary material used for iodine extraction, the functions and regulatory mechanisms of these species have not been fully documented. In this study, a functional genomics analysis of the algal vanadium-dependent haloperoxidase (*vHPO*) gene family was conducted; there genes can introduce halogen atoms into organic compounds. The comprehensive analyses regarding the bioinformatics and phylogenetics of novel genomic and transcriptomic sequencing data of 21 Rhodophyta and 19 Ochrophyta marine algal species revealed that brown algal *vHPOs* have two gene types, vanadium-dependent bromoperoxidase (*vBPO*) and vanadium-dependent iodoperoxidase (*vIPO*), with secondary endosymbiotic host origin. The enzyme activity of *S. japonica* vBPO and vIPO were verified for the first time and were quite stable in a wide range of temperature and pH values. However, the specific activity and optimal conditions were considerably different between vBPO and vIPO. The transcript expression analysis in different *S. japonica* tissues (including rhizoids), generations (sporophytes and gametophytes), sexes (male and female), and stress conditions (hyposaline and hyperthermia) also showed great differences between *vBPOs* and *vIPOs*. Most of the *vBPOs* were constitutively expressed with higher expression dose, which may be responsible for basal halogen metabolism. On the contrary, *vIPOs* mainly showed specific expression, which may be involved in tissue differentiation, generation differentiation, sex differentiation, and stress regulation. Comprehensive analysis of gene family evolution, enzyme biochemical characteristics, and complex transcriptional mechanisms were conducive to the environmental adaptation and sophisticated system evolution of Laminariales. The successful bromination of small-molecule compound substrate by SjavBPO provided high activity and efficient enzymatic tools for artificial synthesis of halogenated compounds.

## Introduction

A halogen substituent is often an essential structural feature of natural products, drugs, or signaling molecules. At present, over 5,000 halogenated compounds have been isolated, including halogenated hydrocarbons, halogenated acetylenes, halogenated phenols, halogenated tyrosine, halogenated fatty acids, and halogenated terpenes (Frank, *et al*., 2016). They are mainly derived through marine algal biosynthesis, which has important biological functions (*e.g*., signaling molecules and defense compounds) and ecological and atmospheric significance (La Barre, *et al*., 2010). Brown algae are widely distributed in temperate and subtropical zones and are an important component of subtidal and intertidal ecosystems (Charrier, *et al*., 2012). The order Laminariales is the most efficient iodine accumulator among all living systems, with an average content of 1.0% dry weight in most *Laminaria* and *Saccharina* species, representing approximately a 30,000-fold accumulation of this element from seawater (Saenko, *et al*., 1978; Leblanc, *et al*., 2006). This is the reason *Laminaria* was previously used in Europe as raw material for iodine extraction (Lüning, 1985) and *Saccharina* in China as a dietary iodine supplement to prevent goiter (Brinkhuis, *et al*., 1987).

Nature has evolved exquisite methods to introduce halogen atoms into organic compounds using halogenating enzymes. The vanadium-dependent haloperoxidase (vHPO) is one of the most studied types among these enzymes due to its biocatalytic properties, including an unusual stability and tolerance for heat and organic solvents (Coupe, *et al*., 2007; Fernández-Fueyo, *et al*., 2015; Sabuzi, *et al*., 2015; Weichold, *et al*., 2016). It can be divided into vanadium-dependent chloroperoxidase (vCPO), vanadium-dependent bromoperoxidase (vBPO), and vanadium-dependent iodoperoxidase (vIPO), depending on the oxidation ability of the halogen ions. The first vHPO to be isolated and characterized was the vBPO from the brown alga *Ascophyllum nodosum* (Vitler, 1984). To date, vHPOs have been characterized from all major classes of marine algae, such as brown algae (Vreeland, *et al*., 1998; Weyand, *et al*., 1999), red algae (Itoh, *et al*., 1986, 1987; Shimonishi, *et al*., 1998; Isupov, *et al*., 2000), green algae (Itoh, 1985; Wever, *et al*., 1985; de Boer, *et al*., 1986; Sheffield, 1993; Ohshiro, 1999; Manley, 2001; Colin, 2003; Suthiphongchai, 2008), as well as terrestrial lichens (Plat, *et al*., 1987), fungi (van Schijndel, *et al*., 1993; Barnett, *et al*., 1998), and cyanobacteria (Frank, *et al*., 2016). These three enzymes, in terms of the gene structure, have the same conservative metal ion binding sites; the histidine residues of imidazole rings, the region for vanadium ions, and amino acid series have a high degree of homology (Littlechild, *et al*., 2002). Overall, *vCPO* gene is mainly found in terrestrial organisms, whereas *vBPO* is mainly found in marine organisms, and *vIPO* is only found in a small number of marine brown algae (Gribble, 2004).

There are few studies on the metabolism of halogens in marine algae. It has been shown that in some brown algae (Wever, *et al*., 1991) and a green macroalga (Manley, 2001), the vBPOs were located at or near the surface of the seaweed. The possible role of the extracellular enzymatic system may be to control the colonization of the surfaces of the seaweed by generating HOBr, which is directly bactericidal (Hansen, *et al*., 2003; Renirie, *et al*., 2008). Until recently, the mechanism of their accumulation, metabolism, and transportation needed further study. In the present study, the first functional *vBPO* and *vIPO* from *S. japonica* were identified. In addition to phylogenetic analysis and transcriptional regulation, this may be worthy of an in-depth study regarding physiological adaptations and relationships with ecological systems and the atmospheric environment. Such research might also provide a platform for diverse protein engineering efforts, and thus an opportunity to establish a new chemoenzymatic halogenation tool in the future.

## Materials and methods

### Algal sample collection

Preserved *S. japonica* haploid gametophytes (male and female) were available as laboratory cultures and were obtained from our Laboratory of Genetics and Breeding of Marine Organisms. Fresh samples of the *Saccharina* sporophytes (rhizoids, stipe, blade tip, blade pleat, blade base, and blade fascia) were collected from east China (Rongcheng, Shandong Province, 37°8′53″N, 122°34′33″E). These samples were used for RNA Sequencing analysis. Various gametophyte samples (male and female) and tissues of sporophytes (rhizoids, stipe, blade tip, blade pleat, blade base, and blade fascia) were collected to analysis relative gene expressions. To detect the influences of abiotic factors, the female gametophytes and blade base of sporophytes were cultured under different temperatures (Control: 8°C; Hyperthermia: 18°C), salinities (Control: 30‰; Hyposaline: 12‰), and circadian rhythms (Control: 30 µmol photons/m^2^·s for 12 h; Darkness: no irradiance for 12 h).

### Sequence analysis

In the present study, genes were identified by analyzing transcriptomic and genomic sequencing data of *S. japonica* (Tao Liu, unpublished data), as well as of the species whose genomes and transcriptomes were sequenced and published in OneKP (www.onekp.com) or NCBI. Matching sequences were manually checked for accuracy with the corresponding known cDNA sequences. The unigenes related to *vHPO* were re-verified using the BLASTX algorithm (http://blast.ncbi.nlm.nih.gov/Blast.cgi). Sequence identities were calculated using the Clustal Omega tool (http://www.ebi.ac.uk/Tools/msa/clustalo/).

### Phylogenetic tree construction

All downloaded sequences are listed in Supplementary Fig. S1. The sequences were aligned using ClustalX 1.83 software (Thompson, *et al*., 1997). The amino acid phylogenetic trees were constructed using MrBayes 3.1.2 software (Ronquist and Huelsenbeck, 2003). The posteriori probability was based on the Metropolis-Hastings-Green algorithm through four chains (Markov Chain Monte Carlo, MCMC) with the temperature set to 0.2°C. The chains were run for 10,000,000 cycles (Posada and Crandall, 1998; Ronquist and Huelsenbeck, 2003). Random trees were constructed in the MCMC analysis, and one tree in every 1,000 generations was saved. After discarding the aging 25% of all tree samples, the residual samples were used to construct a consensus tree; the tree was rendered using Tree View v.1.6.5 software (Page, 1996).

### Purification of recombinant proteins expressed in Escherichia coli

One *vBPO* (*SjavBPO*) and one *vIPO* (*SjavIPO*) from *S. japonica* were synthesized (Shanghai Xuguan Biotechnological Development Co. LTD) to construct recombinant plasmids (NCBI accession number MG195954 and MG195955). *SjavBPO* and *SjavIPO* were cloned in pET32a. These recombinant plasmids were transformed into *E. coli* BL21 (DE3) cells. Isopropyl β-D-1-thiogalactopyranoside (IPTG) was added at a concentration of 0.5 mM to induce the over-expression of the target proteins, and the bacterial cultures were incubated for 16 h at 20°C. His•Bind Resin and GST•Bind Resin were used according to the manufacturer’s instructions (www.yuekebio.com). The proteins were stored at −80°C.

### Assays for enzyme kinetics

The vHPO activity of the purified enzymes was detected using previously described methods (Coupe, *et al*., 2007). For enzymatic characterization, Cl^−^, Br^−^, and I^−^, which are considered potential substrates, were tested. The effect of pH on the enzymatic activities of the purified proteins was determined in the range of 6.0 to 10.0 for SjavBPO1 and 2.5 to 6.5 for SjavIPO1. The effect of temperature on these enzymes was determined over a range of 10°C to 60°C. Four replicates were analyzed for each condition to ensure the consistency of the experimental results. In each case, boiled purified recombinant enzymes were used as the negative control. All data were subjected to a one-way analysis of variance (one-way ANOVA) followed by a Student’s *t*-test.

The small molecular compound HD-ZWM-163 used in the halogen addition experiment was provided by Professor Zhu Weiming from Ocean University of China. It is a type of alkaloid staurosporine extracted from actinomycetes *Streptomyces fradiae* 007 var. M315. Its molecular weight is 466 g/mol, with a purity of more than 98% (Supplementary Fig. S2). HD-ZWM-163 replaced MCD in the above enzyme assay at 20°C for 12 h. The reactants were extracted three times using ethyl acetate and were dissolved in chromatographic methanol. The HPLC-UV test was performed with 75% methanol (v/v, 1.5‰ TFA; flow rate: 1 mL/min), and the peak samples were collected for the determination of the HPLC-MS.

### Transcriptome sequencing

Total RNA was extracted using an improved CTAB method (Gareth, *et al*., 2006). A total amount of 3 µg RNA per sample was used as input material for the RNA sample preparations. Sequencing libraries were generated using NEBNext® Ultra™ RNA Library Prep Kit for Illumina® (NEB, USA) following the manufacturer’s recommendations, and index codes were added to attribute sequences to each sample. The clustering of the index-coded samples was performed on a cBot Cluster Generation System using TruSeq PE Cluster Kit v3-cBot-HS (Illumina) according to the manufacturer’s instructions. After cluster generation, the library preparations were sequenced on an Illumina Hiseq platform, and 125 bp/150 bp paired-end reads were generated. HTSeq v0.6.1 was used to count the reads numbers mapped to each gene. The FPKM of each gene was then calculated based on the length of the gene and the reads count mapped to this gene.

## Results

### Phylogenetic analysis of algal vHPO genes

Novel transcriptomic sequencing data were obtained for 21 Rhodophyta and 19 Ochrophyta marine algal species (OneKP database), and re-sequencing genomic data for *S. japonica* were used to identify *vHPO* genes. Additionally, 104 new full-length candidate genes from 10 brown algal species and 18 genes from 12 red algal species were detected (Supplementary Fig. S1). There were 21 *vBPOs* and 68 *vIPOs* isolated from genomic and transcriptomic data for *S. japonica*. The homology comparison and structure prediction confirmed that these sequences all belonged to the vHPO superfamily.

Phylogenetic trees based on full-length amino acid sequences of *vHPO*-related genes from archaeal taxa, bacteria, fungi, and eukaryotic algae were constructed using Bayesian methods (only representative candidates were included). Based on this tree, all *vHPO* genes formed a monophyletic group sharing a common ancestor with the *vCPO* genes in fungi, after which they evolved independently in red and brown algae (Fig. 1). Red algae only have *vBPO* genes. However, brown algae contain *vBPOs* and *vIPOs*, which both have secondary endosymbiotic host origins. These two types of *vHPOs* were paralogues resulting from an ancestral gene duplication. Interestingly, brown alga *Ectocarpus siliculosus* has only one *vBPO* gene in its genome, whereas the closely related species *S. japonica* has 89 *vHPOs*. Based on the consensus tree, 21 *S. japonica vBPOs* were clustered into three groups (I–III), and 68 *vIPOs* were clustered into five groups (I–V). The large number of family members are expected to have been derived from recent tandem duplication events, which occurred after the differentiation of *Saccharina* and *Ectocarpus*.

**Figure 1.**
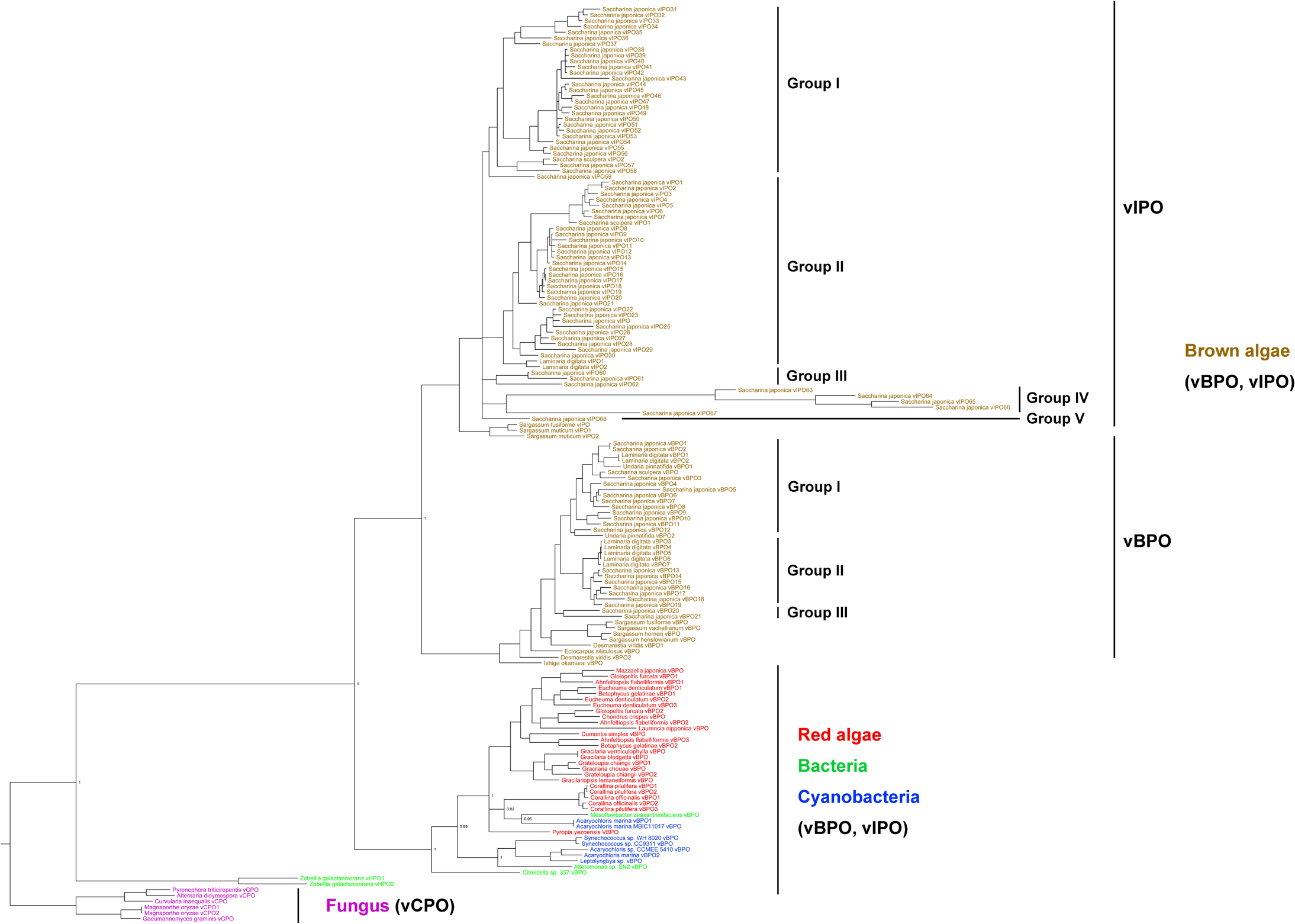
Bayesian phylogenetic tree based on the translated amino acids of *vHPO* with bootstrap values (when >50%) indicated at the nodes. Brown algae *vHPOs* originated from eukaryotic hosts, and underwent gene duplication in their common ancestor. Brown algal vBPOs were clustered into 3 Group (I, II and III), and vIPOs were clustered into 5 Group (I to V). All vHPO sequences were obtained from GenBank or OneKP databases (Supplementary Fig. S1).

### *Characterization and confirmation of the functions of* vHPO *genes from* S. japonica

In this study, one *vBPO* (*SjavBPO1*) and one *vIPO* (*SjavIPO1*) were chosen to be over-expressed in *E. coli* to verify their encoding enzyme activities. This was the first functional analysis of these enzymes in *Saccharina*. Specificity of SjavHPOs was determined by assaying its activity in the presence of different potential substrates. The oxidation ability of halogen ions Cl^−^, Br^−^, and I^−^ were tested. These experiments demonstrated that the SjavBPO1 enzyme exhibited oxidation activity for both Br^−^ and I^−^, whereas SjavIPO1 could only oxidize I^−^ (Table 1). Purified SjavBPO1 had a specific activity of 5861.1 U/mg and 2342.88 U/mg for Br^−^ and I^−^, respectively. SjavIPO1 had a specific activity of 333.6 U/mg for I^−^. These activities were almost the highest of those measured for algal vHPOs (Table 1). In addition, over 90% of the enzymatic activity was detected in SjavHPOs after having been stored at 4°C for 72 h, suggesting that the recombinant proteins were stable under the purification conditions tested.

**Table 1.** The protein activity of vHPOs in different organisms.

| Gene | Species | Iron | Specific activity<br>(U/mg) | Reference |
| --- | --- | --- | --- | --- |
| *SjavBPO* | *Saccharina japonica* | Br<sup>-</sup> | 5861.1 | This study |
| *SjavBPO* | *Saccharina japonica* | I<sup>-</sup> | 2342.9 | This study |
| *SjavBPO* | *Saccharina japonica* | I<sup>-</sup> | 333.6 | This study |
| *vBPO* | *Corallina officinalis* | Br<sup>-</sup> | 9.9-469 | Zhang, *et al.*, 2011 |
| *vBPO* | *Corallina officinalis* | Br<sup>-</sup> | 450-5000 | Coupe, *et al.*, 2007 |
| *vBPO* | *Corallina pilulifera* | Br<sup>-</sup> | 2.32-286 | Shimonishi, *et al.*, 1998 |
| *vBPO* | *Ascophyllum. nodosum* | Br<sup>-</sup> | 50-250 | Hartung, *et al.*, 2008 |
| *vBPO* | *Laminaria digitata* | Br<sup>-</sup> | 42 | Colin, *et al.*, 2003 |
| *vBPO* | *Laminaria digitata* | I<sup>-</sup> | 62 | Colin, *et al.*, 2003 |
| *vIPO* | *Laminaria digitata* | I<sup>-</sup> | 310 | Colin, *et al.*, 2003 |

The activities of the two purified proteins under different temperatures and pH values were determined to elucidate their biological characteristics. Based on these experiments, the maximum activity of SjavBPO1 for Br^−^ was at 20°C, whereas the activity was 68% and 93% of the maximum activity at 10°C and 25°C, respectively. The optimum temperature for I^−^ was 25°C, with 78% and 72% of the residual activity at 20°C and 30°C, respectively (Fig. 2A). For SjavIPO1, the optimum temperature was much higher (50°C), with 68% and 81% of the residual activity at 40°C and 60°C, respectively (Fig. 2B). The optimum pH for SjavBPO1 for both Br^−^ and I^−^ was 6.5, with the activity being 71% to 94% of the maximum activity at other pH values (6.0, 7.0, 8.0, 9.0, 10.0) (Fig. 2C). The optimum pH for SjavIPO1 was determined to be 3.0, and 58% to 99% of the activity remained intact at pH values from 2.5 to 6.5 (Fig. 2D).

**Figure 2.**
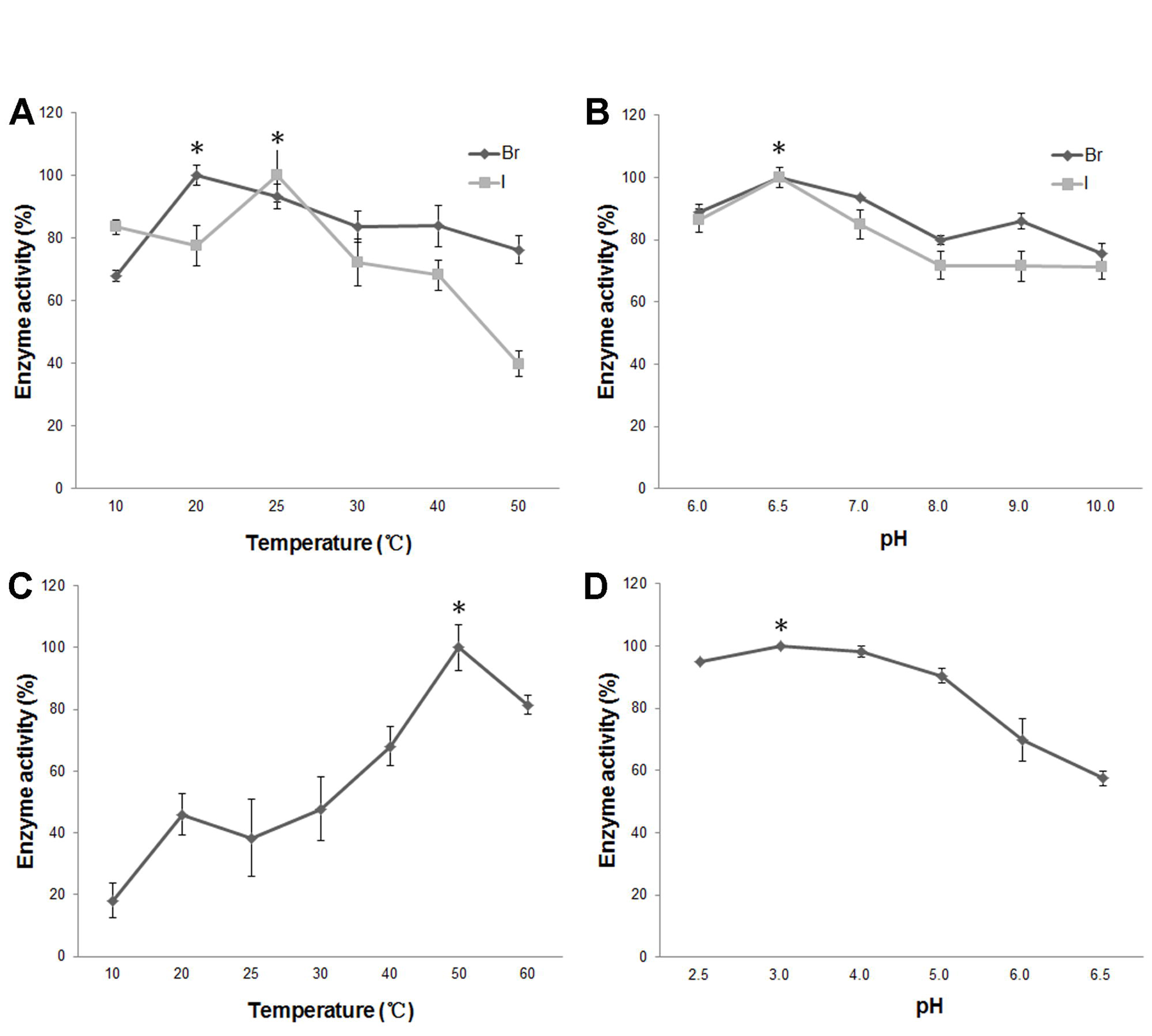
Effects of temperature and pH on activity of SjavBPO1 and SjavIPO1. A. Enzyme activity of SjavBPO1 at 20°C for Br^−^ and 25°C for I^−^ were set to 100%, respectively. B. Enzyme activity of SjavBPO1 at pH 6.5 for both Br^−^ and I^−^ were set to 100%. C. Enzyme activity of SjavIPO1 at 50°C for I^−^ was set to 100%. D. Enzyme activity of SjavIPO1 at pH 3.0 for I^−^ was set to 100%. Data represent means ± SD of four independent experiments.

### Halogen addition of SjavBPO to small-molecule compound

A small-molecule compound (HD-ZWM-163) was used to validate the halogen addition activity of SjavBPO1. In Fig. 3A, a peak of a single compound appears, which is the unmodified compound HD-ZWM-163 (MW = 466 g/mol). The first peak in Fig. 3B and 3C was 2-(N-Morpholino)ethanesulfonic acid (MES) in the reaction Buffer, and peak A and peak B were the derivatives of the halogen addition reaction. Peak B2 was approximately four times that of B1, confirming the effect of the SjavBPO1 protein on the HD-ZWM-163 enzymatic reaction. The samples of peaks A1, A2, B1, and B2 were then detected by mass spectrometry analysis. Compounds A1 and A2 were identified as monobromo-HD-ZWM-163 products (Supplementary Fig. S3), whereas B1 and B2 were dibromo-HD-ZWM-163 products (MW = 623 g/mol, Fig. 3D). In addition, the abundance ratio of peak 623:625:627 is approximately 1:2:1 and also confirms that compound B contain two Br atoms. After adding the SjavBPO1 protein, the production of B2 was more obvious than that of B1, indicating that SjavBPO1 played a role in the compound HD-ZWM-163 dibromination reaction.

**Figure 3.**
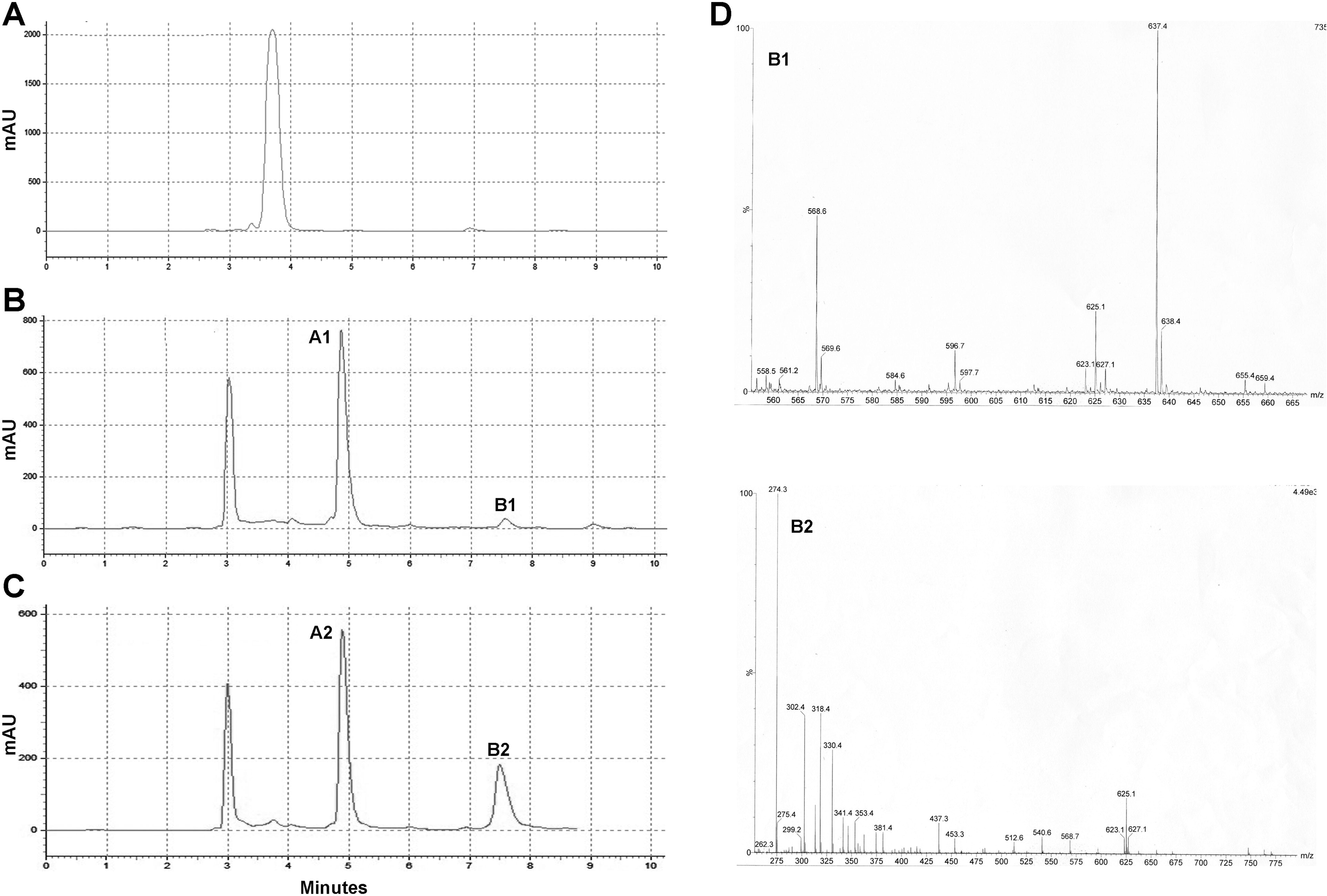
Halogen addition of SjavBPO1 to HD-ZWM-163. (A) The HPLC result of HD-ZWM-163 (flow rate: 1 mL/min; tR = 3.70 min). (B) The HPLC result of HD-ZWM-163 in reaction buffer without SjavBPO1 (tA1 = 4.91 min, tB1 = 7.51 min). (C) The HPLC result of HD-ZWM-163 in reaction buffer with SjavBPO1 (tA2 = 4.91 min, tB2 = 7.51 min). Peaks A1, B1, A2, and B2 were the derivatives of the halogen addition reaction. (D) The ESI-MS result of B1 in (B) and B2 in (C). Compounds B1 and B2 were dibromo-HD-ZWM-163 products.

### Expression differences of Saccharina vHPOs

The expression of 89 *S. japonica vHPO* genes was determined in different generations (sporophytes and gametophytes), tissue (rhizoids, stipe, blade tip, blade pleat, blade base, and blade fascia), sexes (male and female gametophytes), and stress conditions (hyperthermia, hyposaline) (Fig. 4). In all normal and stress conditions, 26.1% (6/23) and 50% (33/66) of *vBPOs* and *vIPOs* were not detected, respectively. However, there were many constitutive *vHPOs* expressed in different tissues (14 *vBPOs* and four *vIPOs*), generations (nine *vBPOs* and eight *vIPOs*), sexes (six *vBPOs* and four *vIPOs*), and stress conditions (seven *vBPOs* and two *vIPOs*). It was determined that *vHPOs* were widely involved throughout the process of *S. japonica* growth, development, and environmental adaptation (Fig. 5).

**Figure 4.**
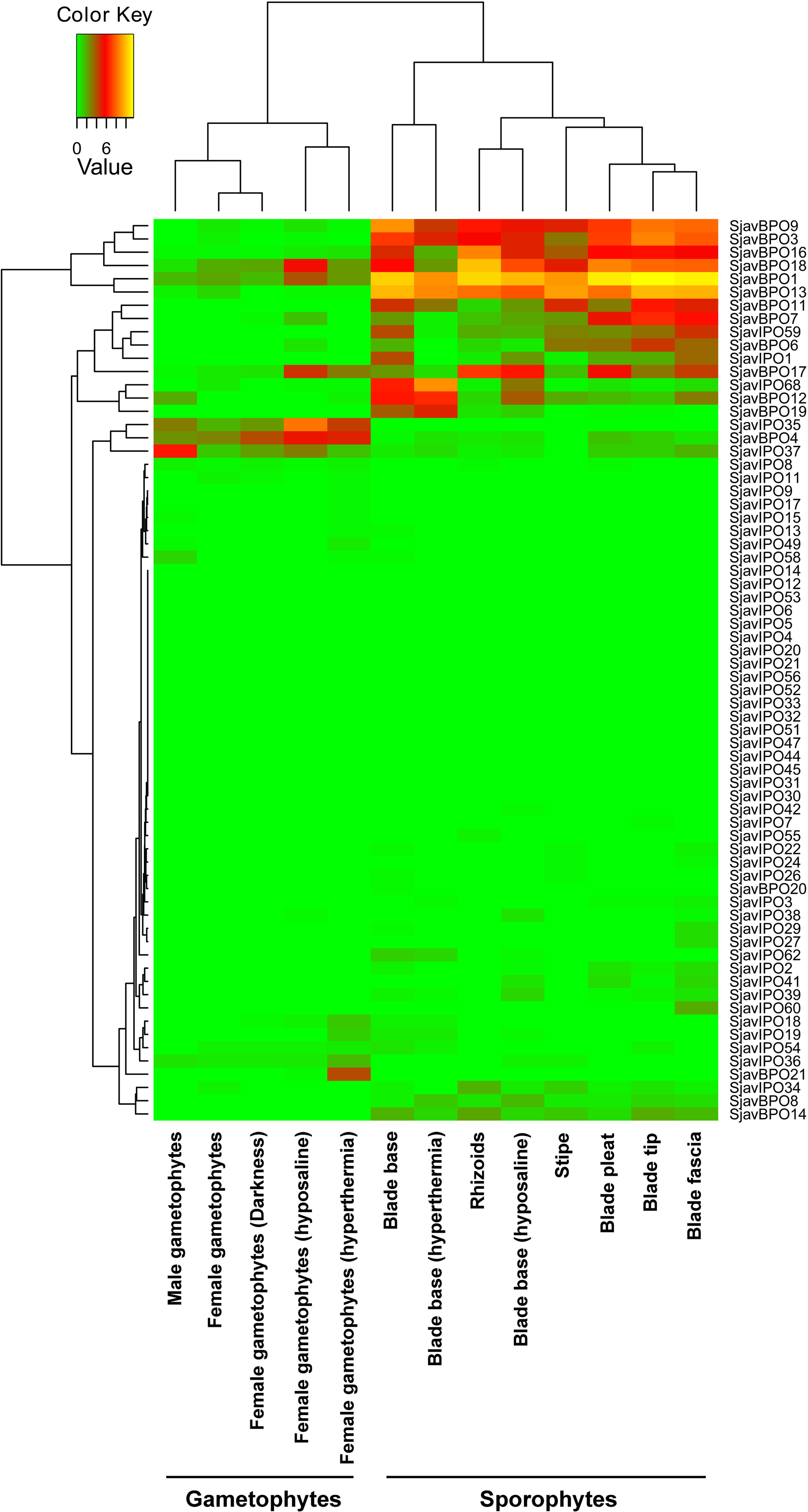
The transcriptional expression of 89 *S. japonica vHPO* genes in different generations (sporophytes and gametophytes), tissue (rhizoids, stipe, blade tip, blade pleat, blade base, and blade fascia), sexes (male and female gametophytes) and stress conditions (hyperthermia, hyposaline).

**Figure 5.**
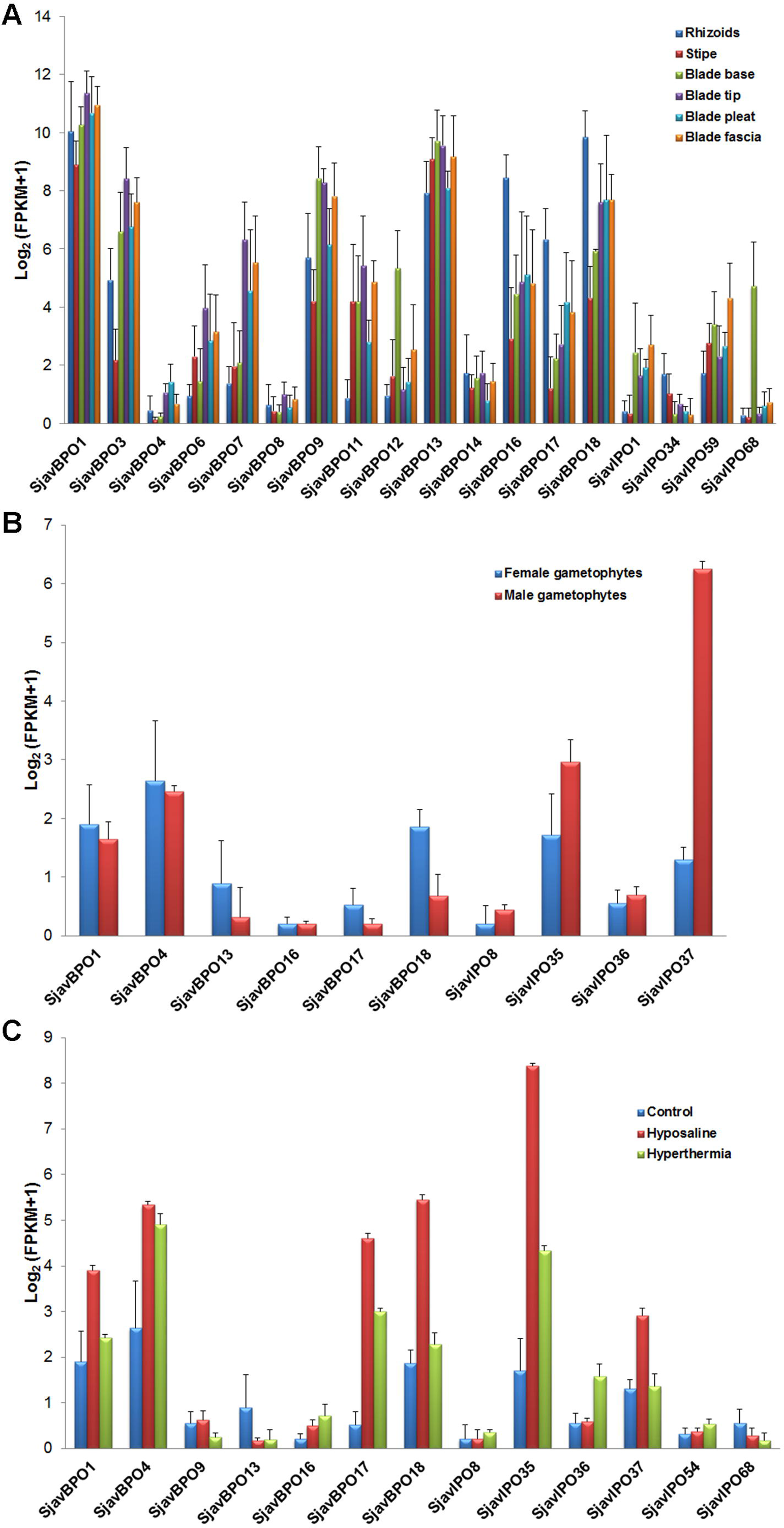
Constitutive expressed *SjavHPOs* in different tissues (A), sexes (B) and stress conditions (C).

The *vBPOs* and *vIPOs* identified in *S. japonica* showed diverse patterns of expression. The expression of *vIPOs* was more specific than was the expression of *vBPOs* (Fig. 6). There were 27 *vIPO* and 16 *vBPO* genes expressed in sporophyte tissues. The specific expression ratio of *vIPOs* (23/27, 85.2%) was 6.8 times higher than that of *vBPOs* (2/16, 12.5%). Sixteen *vBPOs* and 30 *vIPOs* were expressed in different generations, and the specific expression ratios of *vIPOs* and *vBPOs* were 73.3% (22/30) and 43.8% (7/16), respectively. There were nine *vBPOs* and 11 *vIPOs* expressed in male and female gametophytes. The specific expression ratios of *vIPOs* and *vBPOs* were 63.6% (7/11) and 33.3% (3/9), respectively. Female gametophytes displayed a similar response under hyperthermia. The specific expression ratio of *vIPOs* (8/15, 53.3%) was higher than that of *vBPOs* (2/9, 22.2%). On the contrary, the gene number of constitutive expressed *vBPOs* was more than that of *vIPOs*, and expressed constitutive *vBPOs* exhibited higher expression levels than *vIPOs* (Fig. 5). For example, the total expression dose of *vBPOs* in different tissues was about 10^2^–10^3^ times (27.1–520.1 times) that of *vIPOs*.

**Figure 6.**
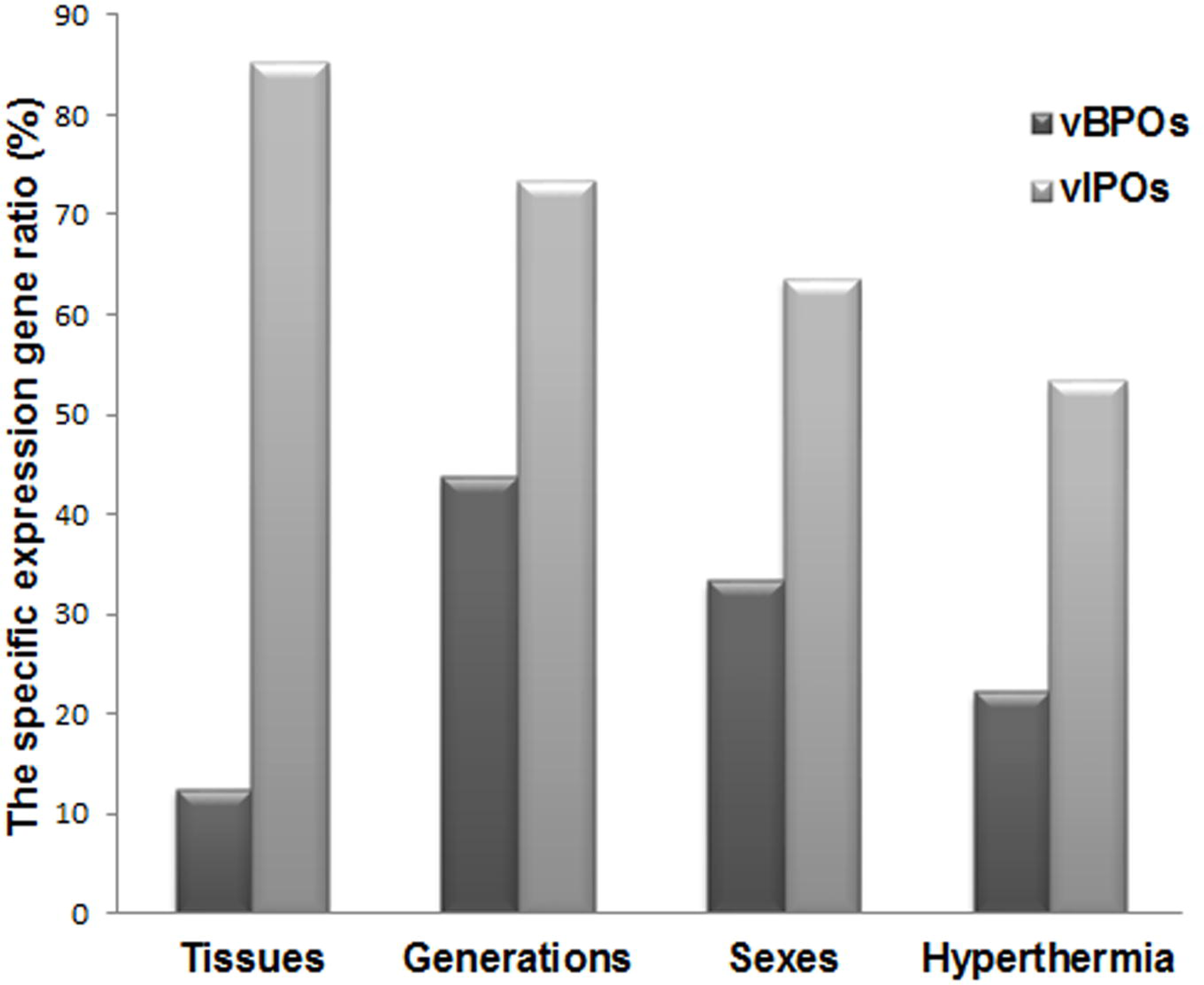
The specific expression gene ratios of *vBPOs* and *vIPOs* in different tissues, generations, sexes and under hyperthermia condition. The expression of *vIPOs* was more specific than that of *vBPOs*.

Considering the different generations, the average expression of sporophyte *vBPOs* (FPKM value = 3307.6) was much higher than that of gametophyte *vBPOs* (FPKM value = 12.7) (P = 0.001), whereas there was no significant difference in *vIPOs* between the two groups (FPKM value = 34.1 vs 45.2) (Fig. 7A). There were 26 *vHPOs* (7 *vBPOs* and 19 *vIPOs*) that were only expressed in sporophytes and three *vIPOs* that were only expressed in gametophytes. Among the sporophytes, some genes were only expressed in unique tissues, such as *vIPO55* in the rhizoids, *vIPO36* in the stipes, *vIPO62* in the blade bases, *vIPO7* in the blade tips, *vIPO11* in the blade pleats, and *vIPO60* in the blade fascia. The expression levels were higher in the blade tips than other sporophyte samples, followed by the blade fascia, rhizoids, blade bases, blade pleats, and stipes (Fig. 7B).

**Figure 7.**
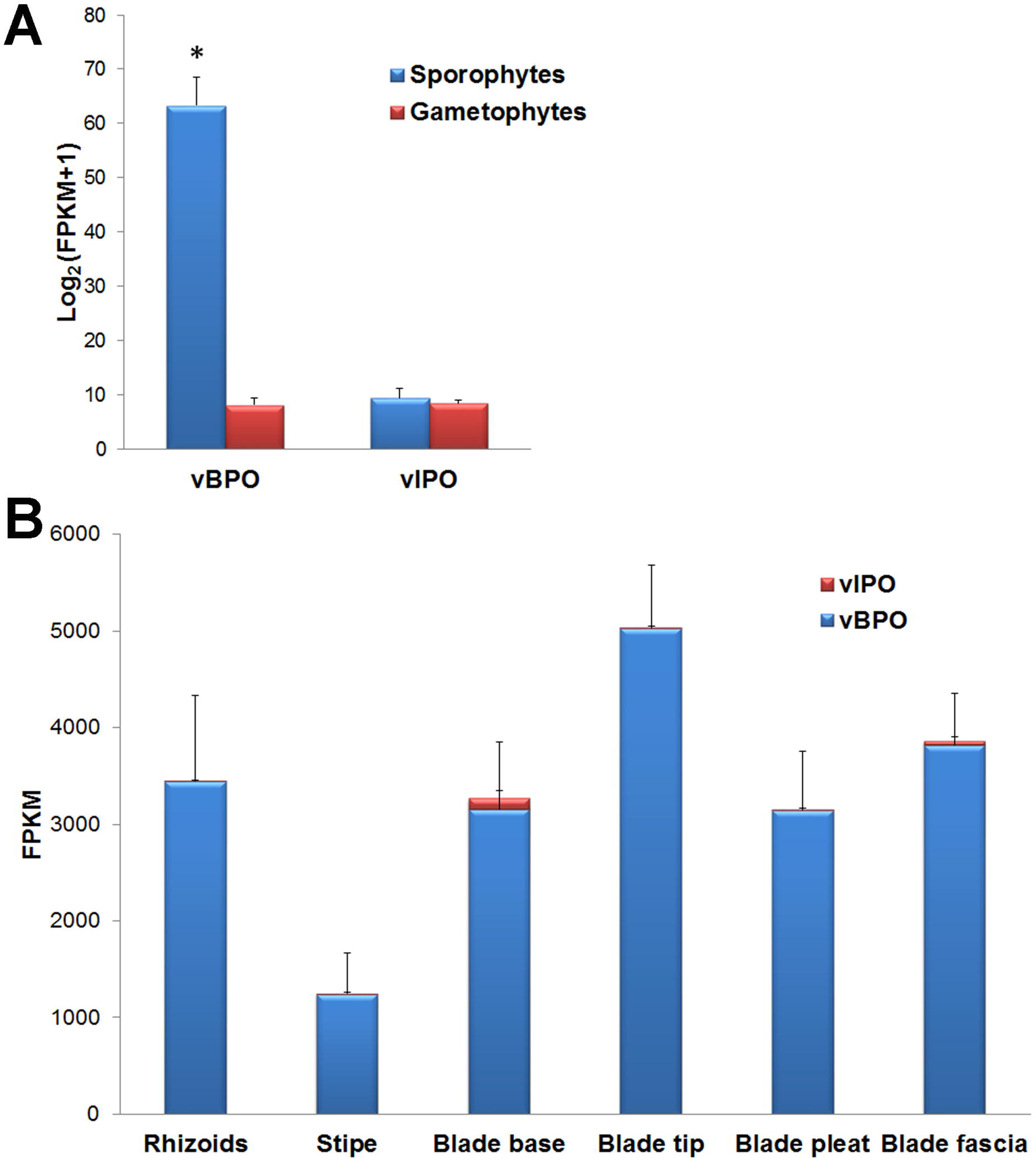
The transcriptional expression of *vBPOs* and *vIPOs* in different sexes (A) and tissues (B). A. The average expression dose of sporophyte *vBPOs* was much higher than that of gametophyte *vBPOs*. The difference between vIPO from sporophyte and gametophyte was not significant. B. The highest total expression dose of *vHPOs* was in blade tip, followed by blade fascia, rhizoids, blade base, blade pleat, and stipe.

The expression of vHPOs differed between male and female gametophytes. Few genes were expressed during the gametophyte stage, including nine *vBPOs* and 11 *vIPOs*. One *vBPO* and three *vIPOs* were specifically expressed in male gametophytes, whereas two *vBPOs* and four *vIPOs* were specifically expressed in female gametophytes. Some of the constitutively expressed genes were highly expressed in female gametophytes (*vBPO18*, P = 0.02), and some were highly expressed in male gametophytes (*vIPO37*, P < 0.01) (Fig. 8).

**Figure 8.**
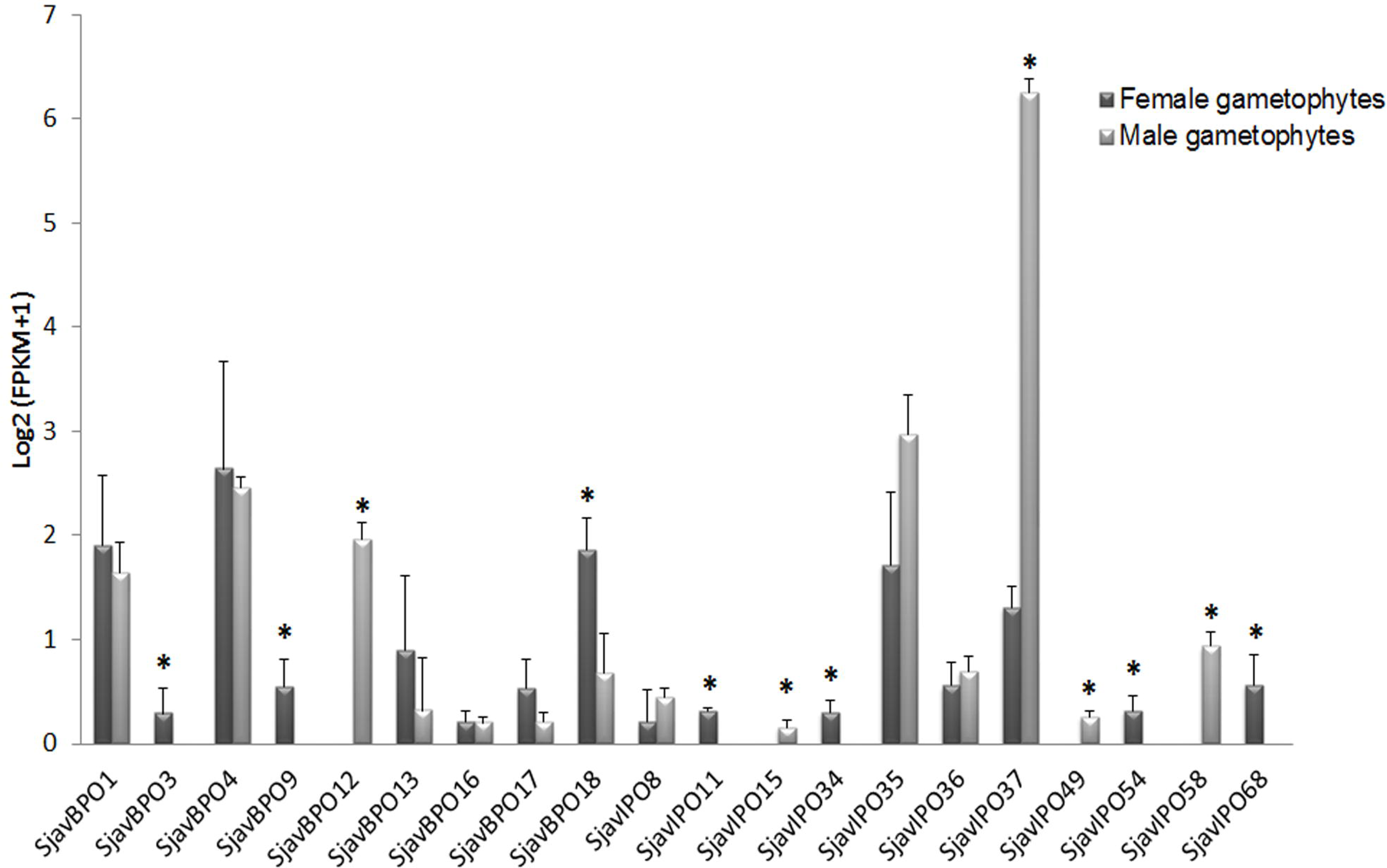
The transcriptional expression of nine *vBPOs* and 11 *vIPOs* in female and male gametophytes. Three *vBPOs* and four *vIPOs* genes were highly expressed in female gametophytes, and one *vBPOs* and four *vIPOs* genes were highly expressed in male gametophytes.

The expression of *vHPOs* under abiotic stress was significantly upregulated in female gametophytes. Compared to gametophyte samples cultured under normal condition, three *vBPOs* (*vBPO6*, *vBPO7*, and *vBPO21*) and three *vIPOs* (*vIPO38*, *vIPO55*, and *vIPO18*) were expressed under hyposaline induction; two *vBPOs* (*vBPO12*, *vBPO21*) and eight *vIPOs* (*vIPO9*, *vIPO13*, *vIPO15*, *vIPO17*, *vIPO18*, *vIPO19*, *vIPO49*, and *vIPO58*) expressed under hyperthermia (Fig. 9). However, the expression of the *vHPOs* in gametophytes did not change significantly under the normal circadian rhythm (12 h irradiation vs. 12 h darkness) or in sporophytes under hyposaline or hyperthermia stresses.

**Figure 9.**
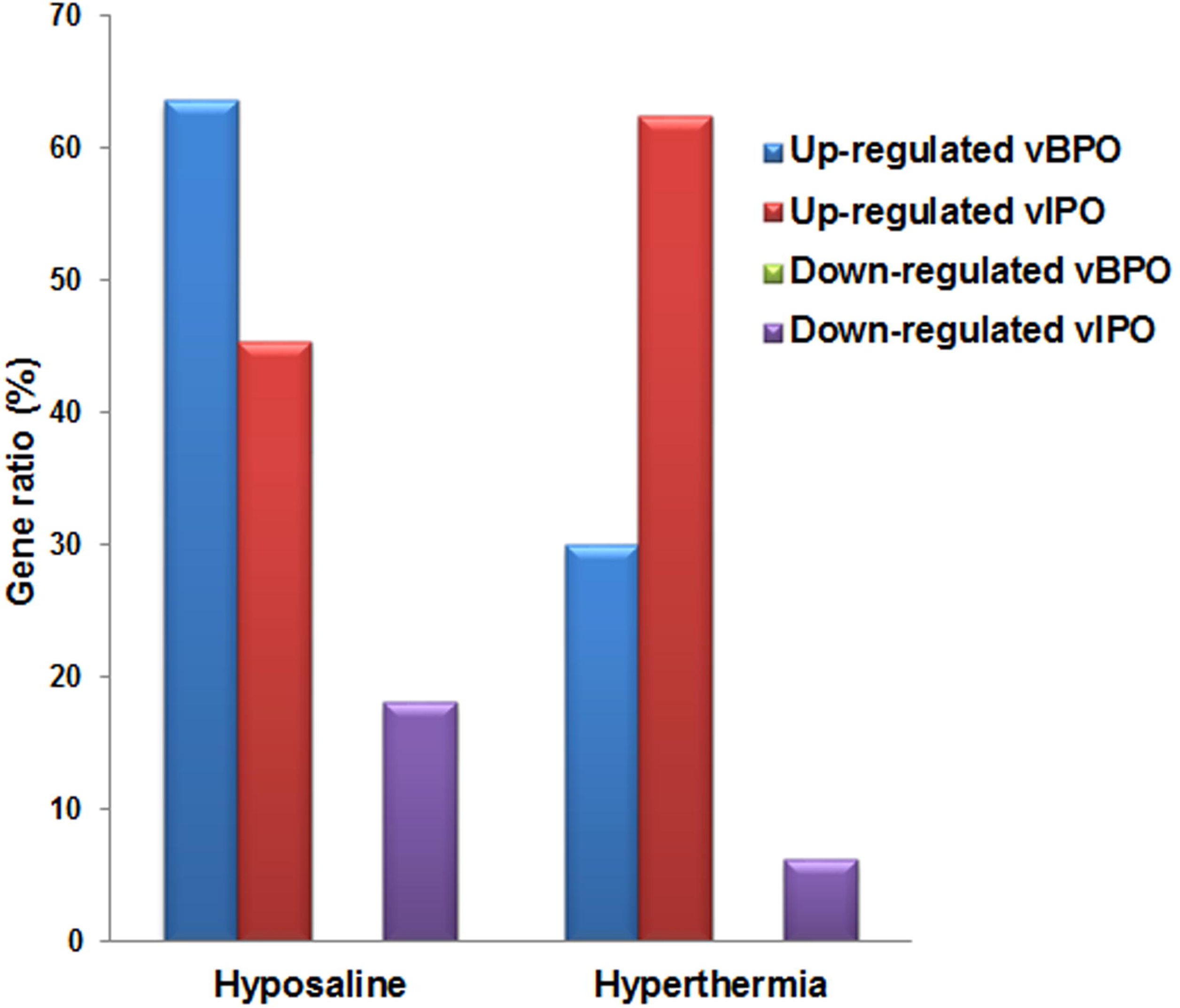
The up-regulated and down-regulated gene ratio of *vBPOs* and *vIPOs* under hyposaline and hyperthermia stresses in female gametophytes.

## Discussion

### *The origin and evolution of* vHPOs *contributed to the differentiation of brown algal* vBPO *and* vIPO *genes*

Our new data allow for the identification of *vHPO* genes widely distributed among different red (such as Gigartinales, Gracilariales, and Halymeniales) and brown (such as Laminariales, Desmarestiales, Ishigeales, and Fucales) algal taxa (Fig. 1). Among the *vHPO* gene family, *vCPO* appeared first in fungi, followed by *vBPO* and *vIPO* with functional specialization. The eukaryotic algal *vHPOs* clustered with the *vCPO* genes of fungi, indicating its endosymbiotic host origin. The differentiation time of *vBPOs* and *vIPOs* could be later than that of the *vCPOs*, which might have occurred because of variations in the earth’s environment. Previous results have demonstrated the independent evolution of *vHPOs* in red algae and brown algae (Ye, *et al*., 2015). This study obtained several *vHPO* genes from several algal species and performed a comprehensive phylogenetic analysis using a wide range of algal species, clarifying their evolutionary relationships.

Furthermore, there are diverse gene duplications among most red and brown algal taxa, as they have undergone gene duplications subsequently at different evolutionary time scales. Because many brown algae were included in the analysis, we confirmed that gene duplications occurred both before (different groups of *SjavBPOs* and *SjavIPOs*) and after species differentiation (such as *SjavBPO1* and *SjavBPO2*, *SjavIPO1* and *SjavIPO2*). This suggested that the *vHPO* gene family in brown algae recently underwent, and perhaps is still undergoing, rapid evolution. Through genomic sequencing analysis, *E. siliculosus* has only one *vBPO*. However, in Laminariales species, the gene family expanded dramatically, especially in the genera *Saccharina* and *Laminaria*. As we know, the iodine content is much higher in these two genera. Vinogradov (1953) demonstrated that the iodine content in *L. digitata* on the west coast of France was as high as 1.7%. Ji (1963) showed that the iodine content in *S. japonica* was also greater than 1%. The appearance of functional *vHPOs* indicated the more efficient immobilization of halogen.

### The diverse enzyme characteristics of SjavBPO1 and SjavIPO1 were conducive to better environmental adaption

The enzyme activities of *S. japonica* vBPO1 and vIPO1 were confirmed (Fig. 2), verifying the authenticity of the pathways as determined through a bioinformatical approach. The oxidation activity of SjavBPO1 is now known as the highest activity of vBPO and vIPO proteins, compared to those measured for algal vHPOs listed in Table 1. This also suggested the massive accumulation of halogen in *S. japonica*. Under the same experimental conditions, the SjavBPO1 protein exhibited no activity with Cl^−^, and the specific activity with Br^−^ was three times that of the activity with I^−^. Meanwhile, the SjavIPO1 protein only had the activity with I^−^. This demonstrated the functional differentiation and specialization between SjavBPO and SjavIPO.

The biochemical characteristics of SjavBPO1 and SjavIPO1 differed considerably. The optimal pH value for SjavBPO1 (pH = 6.5) was similar to that of brown alga *L. digitata* vBPO and vIPO (optimal pH = 5.5; Colin, *et al*., 2003) and red alga *Corallina officinalis* vBPO (optimal pH = 7.0–7.5; Coupe, *et al*., 2007); SjavIPO1 activity was lower (optimal pH = 3.0) (Fig. 2). In contrast, the SjavIPO1 protein was active up to 60°C, which was the optimal temperature for maximum activity, similar to *L. digitata* vIPO (optimal temperature = 50°C; Colin, *et al*., 2003) and *C. officinalis* vBPO (optimal temperature 65°C–70°C; Coupe, *et al*., 2007). The SjavBPO1 maximum activity was considerably lower at 20°C and 25°C for Br^−^ and I^−^, respectively; this is similar to the optimal temperature for *L. digitata* vBPO (30°C) for I^−^ (Colin, *et al*., 2003). The variation of optimum conditions for different proteins indicates that they may play a role under different environmental conditions. Additionally, more than 40% of the maximum activity remained intact in the temperature range of 20°C–50°C for both SjavBPO1 and SjavIPO1 enzymes. For pH stability, over 70% of the maximum activity was in the range of 6.0–10.0 for SjavBPO1, and over 57% of the maximum activity was in the range of 2.5–6.5 for SjavIPO1. These results suggested the high stability of SjavHPO enzymes, which remain active under a wide range of variable conditions. The iodine content of kelp increases with seawater depth (Saenko, *et al*., 1978) and differs between growing seasons (Ar Gall, *et al*., 2004). In the present study, only two representative SjavHPOs were analyzed. The dozens of remaining proteins in *S. japonica* might exhibit different biochemical characteristics. These characteristics could help this species to use halogen and facilitate signal transduction and self-defense under different environmental conditions.

### The complex regulation of vHPOs was a driving force for the sophisticated system evolution in brown algae

The *SjavHPO* gene expression showed significant differences in various tissues, during different developmental stages, and under different types of stress (Fig. 4). Firstly, a large number of genes were not transcribed in the tested samples, which may be redundant gene backup, or were only expressed under special conditions. In contrast, the constitutively expressed genes maintained basic halogen metabolism under different conditions (Fig. 5). Halogen compounds have important ecological and physiological value for algae, such as free-oxygen cleaning and antimicrobial defense (Renirie, *et al*., 2008; Hansen, *et al*., 2003). They form an important source of bromine to the troposphere and lower stratosphere, and they contribute significantly to the global budget of halogenated hydrocarbons (Wever and van der Horst 2013). Additionally, constitutively expressed vHPOs have great physiological and ecological significance.

The expression of *vIPOs* was more specific than that of *vBPOs* in different tissues, generations, sexes, and under hyperthermia stress (Fig. 6). However, the expression doses of *vBPOs* were far higher than those of *vIPOs* in sporophytes (Fig. 4). With respect to the higher specific activity of SjavBPO1 than that of SjavIPO1 (Table 1), *vBPOs* were supposed to be the core of *S. japonica* halogen metabolism, and responsible for the basal halogen accumulation. In contrast, *vIPOs* are mainly involved in generation differentiation, tissue differentiation, sex differentiation and stress regulation.

*S. japonica* possess heteromorphic haploid-diploid life cycles with a macroscopic thallus sporophyte and microscopic gametophyte generation. The gametophyte is similar to some filamentous brown algae, with an isomorphic haploid-diploid filamentous generation (Cock, *et al*., 2014; Bartsch, *et al*., 2008). Therefore, the comparison between *S. japonica* sporophytes and gametophytes might provide an explanation for the evolution of brown algae from unicellular to multicellular organisms (Chi, *et al*., 2017). The expression level was higher in sporophytes than in gametophytes (Fig. 7A). This differs from previous studies (Ye, *et al*., 2015), likely because we annotated more *Saccharina vHPO* genes (89 *vHPOs* in our study vs. 76 *vHPOs* in Ye, *et al*., 2015), and the materials were derived from different *Saccharina* strains. The higher expression in the blade tips (Fig. 7B) is consistent with the higher content of iodine elements in Laminariales distal blades (Shaw, 1962; Ar Gall, *et al*., 2004; Küpper, *et al*., 2008). The blade tips are more sensitive to environmental stresses and pathogen infections than are the regions beneath the thalli (Küpper, *et al*., 1998). This increased gene expression led to greater halogen accumulation. The reserved material might move toward the halogen-requiring basal meristems through the highly specialized elongated sieve elements (Amat and Srivastava, 1985). The differences in gene expression and synthetic product translocation are conducive to the functional differentiation of tissues and necessary for the supply of halo-containing compounds in brown macroalgae. In addition, *SjavHPO* expression was significantly upregulated in the gametophytes under stress, such as hyperthermia and hyposaline conditions (Fig. 9), indicating that *vHPOs* present in *S. japonica* were involved in stress resistance. However, the expression level in sporophytes did not change under the same abiotic stresses. One possible explanation is that the gametophyte stage is more vulnerable to external stress than is the sporophyte stage.

### Algal vHPOs have important industrial and pharmaceutical values

Halo-containing compounds, such as acetogenins (anti-microbial activity), and indoles (anti-inflammatory and anti-cancer activities) could have medical applications (Butler, 1998). However, conventional chemical manufactures generate potentially harmful byproducts. For example, synthetic bromination typically yields approximately 50% of the remaining bromine in the form of waste compounds. Currently, biohalogenation approaches are particularly valuable as an alternative, which could markedly reduce the amount of halogen pollutants produced. For example, using a biotransformation approach employing haloperoxidases to produce drugs such as Vancomycin, Maracen A, and cryptophycins (all halogen-containing compounds) could markedly reduce toxic levels in wastewater (Coupe, *et al*., 2007). Therefore, the halogenation of SjavBPO on small molecule compounds (Fig. 3) could provide a more efficient and convenient method for biohalogenation, which has important economic value and environmental significance.

In conclusion, the huge diversities in gene family members, enzyme catalytic activities, biochemical characteristics, and gene expression patterns of brown algal *vBPOs* and *vIPOs* exhibited great differences between large individual parenchyma (sporophyte, 2n) and single-row filamentous cells (gametophyte, n) regarding biochemistry, physiology, and ecology. The Phaeophyceae-exclusive vIPOs, with a large number of gene expansions and differential expression, and function in scavenging free oxygen, play a crucial role in the evolution of unicellular organisms (such as filamentous *S. japonica*) into multicellular organisms (such as *S. japonica* thalli). The deep resolution of the *vHPO* gene family in brown algae has important biological, ecological, and economic significance.

## Supplementary data

Supplemental Figure S1. List of vHPO sequences used for phylogenetic analysis.

Supplemental Figure S2. The chemical structure of small-compound HD-ZWM-163.

Supplemental Figure S3. The ESI-MS result of peak A1 in Figure 3B and peak A2 in Figure 3C.

## Acknowledgements

This work was supported by the National Natural Science Foundation of China (NSFC No. 41376143), Leading Talents Program in Taishan Industry of Shandong Province, Leading Talents Program in Entrepreneurship and Innovation of Qingdao, China-ASEAN Maritime Cooperation Fund “China-ASEAN Center for Joint Research and Promotion of Marine Aquaculture Technology”, and China Agriculture Research System (CARS-50).

